# Dietary lipids, not ketone body metabolites, influence intestinal tumorigenesis in a ketogenic diet

**DOI:** 10.1101/2025.06.06.658169

**Authors:** Jessica E.S. Shay, Fangtao Chi, Constantine N. Tzouanas, Johanna Ten Hoeve, Kevin J. Williams, Tolga Sever, Isabela Fuentes, Ömer H. Yilmaz

## Abstract

Diet composition shapes tissue function and disease risk by modulating nutrient availability, metabolic state, and cellular dynamics. In the gastrointestinal tract, obesogenic high-fat diets enhance intestinal stem cell activity and tumorigenesis. However, the impact of ketogenic diets (KD), which contain even higher lipid content but induce ketogenesis, remains poorly understood. This is particularly relevant for patients with familial adenomatous polyposis (FAP), who face a high risk of small intestinal tumours. Here, we combine dietary, genetic, and metabolic manipulations in mouse models of spontaneous intestinal adenoma formation to dissect the role of systemic and epithelial ketogenesis in intestinal cancer. We show that KD accelerates tumour burden and shortens survival, independent of ketone body production. Through genetic manipulation of the ketogenic pathway, we modulate local and systemic ketone body production; however, neither inhibition nor augmentation of the ketogenic enzyme HMGCS2 nor disruption of ketolysis altered tumour progression. In contrast, inhibition of fatty acid oxidation did limit adenomatous formation. These findings reveal that dietary lipid content, through FAO rather than ketone body metabolism, influences intestinal tumorigenesis and highlight the need for nuanced consideration of dietary strategies for cancer prevention in genetically susceptible populations.

## Introduction

The influence of diet on tissue homeostasis and cancer risk is a growing concern ^1^. The intestinal epithelium is among the most metabolically dynamic tissues in the body, renewing every 4-5 days under homeostatic conditions ^2^. This regenerative process is orchestrated, in large part, through Lgr5⁺ intestinal stem cells (ISCs) residing at the base of the small intestinal and colonic crypts ^3,4^. Nutrient availability and dietary composition are increasingly recognized as critical modulators of ISC behavior, with growing evidence indicating ISC metabolism influences both cell fate and tumorigenic potential ^5,6^. Obesogenic high-fat diets (HFD), emblematic of Western dietary patterns, have been shown to exert pleiotropic and tumorigenic effects throughout the intestinal epithelium ^7–10^. HFD promotes the expansion of Lgr5⁺ ISCs, enhances their clonogenic potential, and increases the likelihood of malignant transformation with additional effects through PPAR-mediated fatty acid oxidation (FAO) and immune evasion ^11–16^.

Dietary interventions like caloric restriction (CR) and short-term 24 hour fasting confer protective effects against age-associated ISC decline and boost stem cell-mediated repair^17–24^. With respect to tumorigenicity, we recently found that post-fast refed ISCs and early progenitors initiate greater numbers of tumours in the small intestine and colon upon loss of the *Apc* tumour suppressor gene compared to the ad libitum or fasted states ^24^. Ketogenic diets (KDs), while distinct from CR and fasting due to high fat and minimal carbohydrate content, similarly prolong healthspan in mice and demonstrate anti-tumorigenic effects in models of colorectal tumorigenesis; yet, their effects on small intestinal tumorigenesis is less understood ^25–27^.

Nutrient availability, metabolic adaptation, and stem cell function are intimately intertwined, with dietary interventions exerting profound effects on tissue homeostasis, regeneration, and tumorigenesis. The effects of KD on tumorigenesis are somewhat mixed. In an acute myeloid leukemia model, KD accelerated disease progression and reduced survival ^28^. Similarly, in BRAF V600E mutant melanoma, elevated acetoacetate ketone metabolite levels associated with KD enhanced tumour growth ^29^. In a pancreatic cancer allograft model, caloric restriction suppressed tumour growth, whereas KD had no effect ^30^. However, KD has demonstrated significant anti-tumorigenic effects in models of colorectal tumorigenesis. Recent work suggests these protective effects are mediated specifically by ketone body metabolites, particularly beta-hydroxybutyrate (BHB), which acts through the hydroxycarboxylic acid receptor 2 (HCAR2) to suppress colorectal cancer development. The therapeutic potential of these findings is further highlighted by ongoing clinical investigation of beta-hydroxybutyrate as a chemopreventive agent in Familial Adenomatous Polyposis (FAP) syndrome (NCT06578637), an autosomal dominant hereditary condition characterized by the development of hundreds to thousands of adenomatous polyps in the colon and rectum with near-certain progression to colorectal cancer if left untreated, as well as an increased risk of small intestinal adenomas and cancer ^31^.

3-hydroxy-3-methylglutaryl-CoA synthase 2 (HMGCS2), the rate-limiting mitochondrial enzyme in ketogenesis, catalyzes the condensation of acetyl-CoA and subsequent downstream production of the ketone bodies acetoacetate and beta-hydroxybutyrate (BHB) ^32^. While the liver is the principal site of systemic ketone body production, HMGCS2 is also highly expressed in small intestinal ISCs. It is further induced by fasting and ketogenic diets, facilitating the local production of ketone bodies that promote ISC maintenance, proliferation, and regenerative responses through mechanisms such as histone deacetylase inhibition and Notch activation ^17,18^. However, in the context of FAP, where germline APC mutations drive ISC-mediated adenoma formation, the consequences of enhanced ISC function are not straightforward. This is particularly concerning given that, despite prophylactic colectomy mitigating CRC risk in FAP, extracolonic malignancies have become the leading source of morbidity, with small intestinal, especially duodenal, adenomas and cancer developing in 90% and 10% of patients, respectively ^33–37^.

Understanding the differential impact of ketogenic diets on small intestinal versus colonic tumorigenesis is therefore warranted, especially for FAP patients. While evidence suggests ketone body metabolites like BHB may suppress colorectal cancer development, their role in the small intestine - where FAP patients face the highest cancer risk after prophylactic colectomy - remains undefined. This knowledge gap is particularly significant as dietary interventions, including ketogenic approaches, are increasingly considered as adjunctive strategies in cancer management ^38^. Thus, a paradox exists between the lipid-driven enhancement of stemness and the proposed ketone-mediated suppression of tumours ^27^. To explore this dichotomy, we combine dietary, genetic, and metabolic manipulations of ketone body metabolite production with inducible Apc loss to determine how modulation of epithelial and systemic ketone body production and utilization impacts tumorigenesis.

## Results

### Ketogenic diet augments ISC function and increases tumorigenesis

To study the effects of ketogenesis and ketone body metabolites on the small intestine, we took advantage of a previously defined and characterized purified 80% lard-based ketogenic diet (KD) (**Fig. 1a, Extended Data Fig. 1a**, **Table 1**). In keeping with prior findings, mice on a KD exhibited modest weight changes (**Fig. 1b, Extended Data Fig**. **1b, c**) and significantly higher circulating ketones compared to mice on a purified control diet (CD) (**Fig. 1c**). Although no significant changes in glucose were noted (**Fig. 1d**), KD mice had decreased circulating insulin when compared to CD mice (**Fig. 1e**). Using the novel Lgr5-2A-EGFP-2A-CreERT2 reporter mouse with GFP expression in all intestinal crypt stem cells, we observed an increase in Lgr5 GFP^+^ crypt base cells and augmented proliferation as seen by BrdU^+^ staining (**Fig. 1f, g**) ^39^. Niche-promoting Paneth cells were decreased (**Extended Data Fig. 1d, e**), indicating that KD ISCs acquire niche independence as observed in an obesogenic HFD model ^11^. Supporting this notion further, *in vitro* organoid-forming ability was significantly increased in mice exposed to KD (**Fig. 1h, i**), and isolated GFP-expressing stem cells exhibited increased *in vitro* clonogenic growth (**Extended Data Fig. 1f-h**). Lastly, a KD also significantly increased tissue beta-hydroxybutyrate (BHB) levels and the expression of key ketogenic enzymes, such as HMGCS2 (**Fig. 1j, Extended Data Fig. 1i**).

**Table 1:**
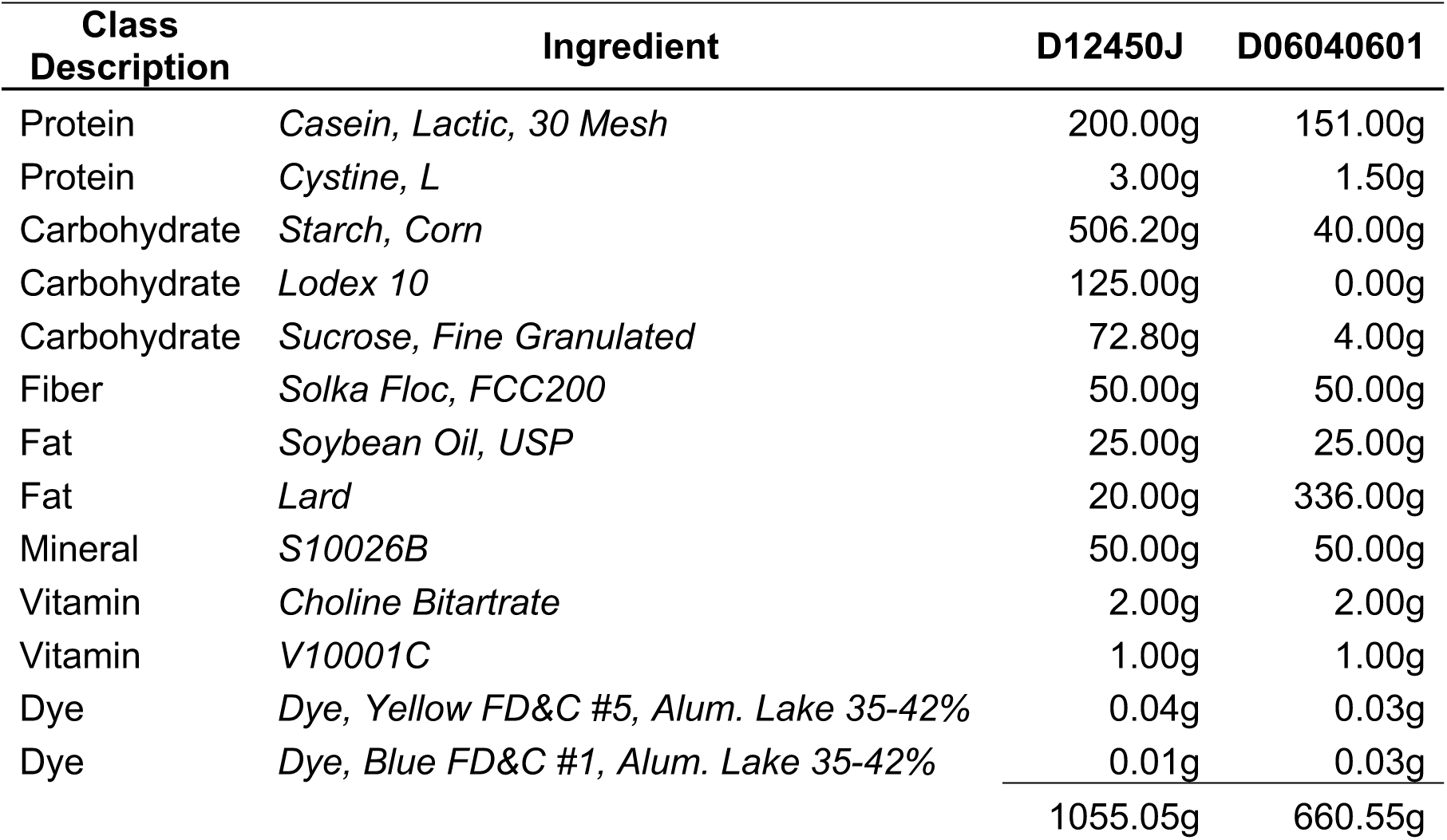

**Figure 1.**
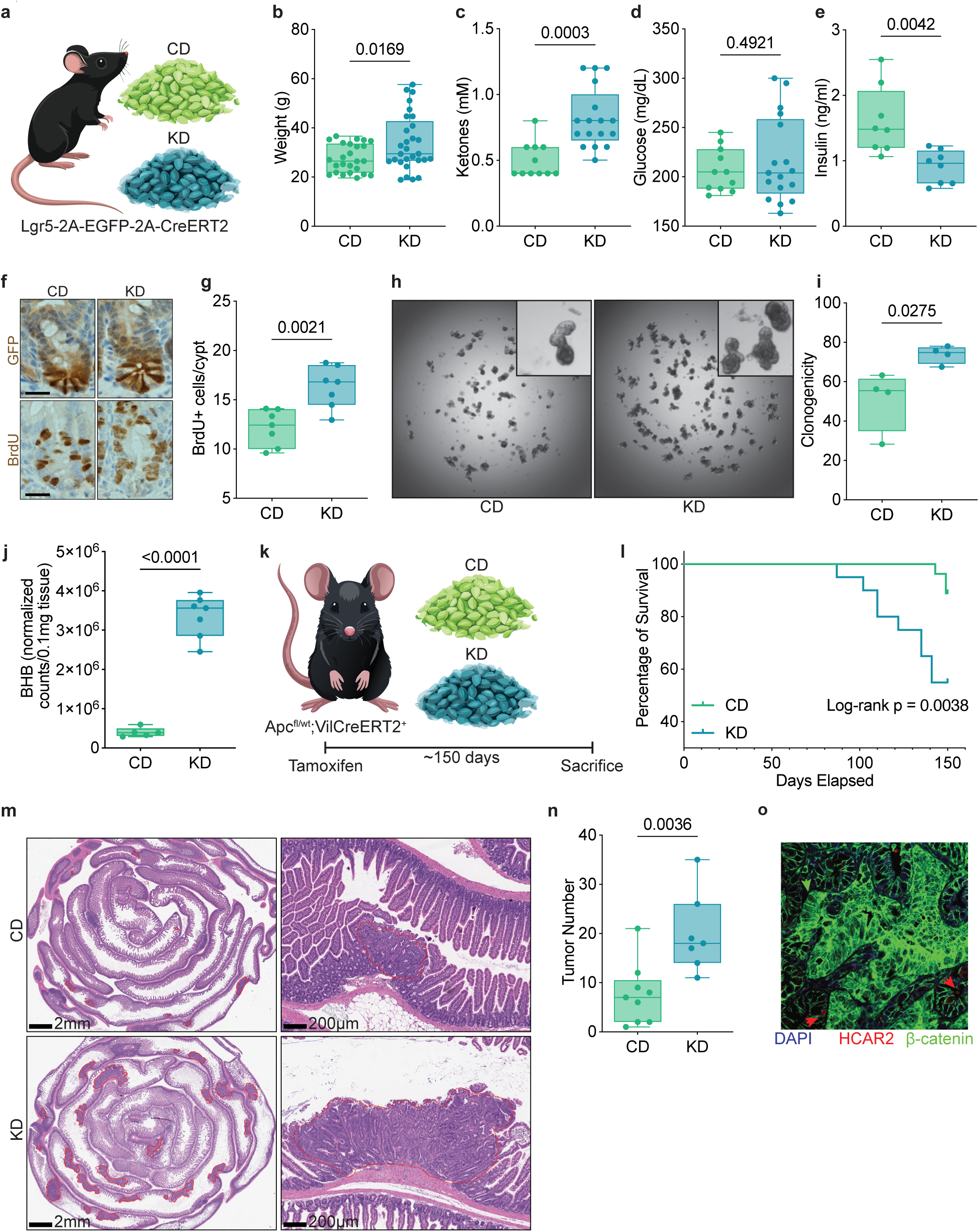
Ketogenic diet enhances intestinal stem cell activity and promotes tumorigenesis. (a) Schematic of experimental design utilizing ketogenic diet (KD) in Lgr5-2A-EGFP-2A-CreERT2 mice. (b) Body weight of control diet (CD) and KD-fed mice over the study period. (c) Circulating β-hydroxybutyrate (BHB) levels in KD-fed mice compared to CD-fed controls. (d) Blood glucose levels in CD- and KD-fed mice. (e) Circulating insulin levels in KD-fed mice relative to CD-fed mice. (f) Representative immunohistochemistry images of Lgr5-GFP+ crypt base cells and BrdU+ cells in CD- and KD-fed mice. (g) Quantification of BrdU+ proliferative cells in the intestinal crypt. (h) Representative images and (i) quantification of *in vitro* organoid-forming assays from crypts isolated from CD- and KD-exposed mice. (j) Intestinal tissue BHB levels in CD- and KD-fed mice. (k) Schematic of tumorigenesis protocol using Vil-CreERT2;Apc^fl/wt^ mice following tamoxifen induction and dietary randomization. (l) Kaplan-Meier survival curves with log-rank Mantel-Cox test comparing CD- and KD-fed cohorts. (m) Representative images and (n) quantification of tumour burden in KD-fed mice relative to CD-fed mice. (o) Immunofluorescence of nuclear β-catenin, HCAR2 and DAPI in early adenomas. Data are mean ± s.d.

To directly assess the impact of dietary lipid content and ketogenesis on intestinal tumorigenesis, we utilized Vil-CreERT2 mice that also carry a single Apc floxed allele, allowing for the timed loss of a single copy of Apc upon exposure to tamoxifen. The spontaneous loss of the second Apc allele via loss of heterozygosity leads to adenoma formation. After receiving the tamoxifen injection, mice were then placed on either CD or KD for approximately 150 days (**Fig. 1k**). During this period, a notable increase in premature death was observed, which correlated with the dietary lipid content. KD-exposed mice have shortened survival curves when compared to CD mice (**Fig. 1l**). Tumour burden was increased in KD (**Fig. 1m, n, Extended Data Fig. 1l-o**). Regardless of diet, a higher tumour burden exhibited a positive association with serum ketone measurements and a negative association with serum glucose levels measured at the time of tissue collection (**Extended Data Fig. 1j, k**). The pro-tumorigenic effect of KD was even more pronounced at 90 days before the survival curves diverged, indicating that tumour burden likely has a direct effect on mouse survival (**Extended Data Fig. 1l-n**). Hydroxycarboxylic acid receptor 2 (HCAR2) has been previously implicated as a BHB receptor that drives the anti-tumorigenic effects of ketone body metabolites in the colon ^27^. Interestingly, HCAR2 expression appeared restricted and was absent from early adenomas where nuclear beta-catenin, a hallmark of APC loss and adenoma development, does not co-localize with HCAR2 (**Fig. 1o, Extended Data Fig. 2a-b**). We sought to examine which cell types expressed Hcar2. In a single-cell RNA sequencing (scRNA-seq) dataset of mouse small intestinal epithelial cells (Aliluev et al; four mice on a high-fat high-sucrose diet, 4205 cells) Hcar2 was detected in 0.2-0.5% of goblet and enteroendocrine cells, respectively, but not in stem cells (**Extended Data Fig. 2c-g**). Hcar2 was also not detected in a separate mouse scRNA-seq intestinal epithelial cell dataset (Mana et al.; eight mice, 62,687 cells), and only a single Hcar2 transcript was detected across 15,053 cells in our scRNA-seq of mice on ketogenic vs. control diets (this study). As such, in the small intestine, a ketogenic diet likely exerts tumour-modulating effects in an HCAR2-independent manner in contrast to what has been proposed in the colon.

### Ketone body metabolites are dispensable for the tumorigenic effects of KD

To further characterize how KD exposure modulates intestinal epithelial biology, we performed scRNAseq on intestinal epithelial cells isolated from control diet (CD) and KD-treated mice. A total of 15,053 cells were captured and analyzed across conditions (**Fig. 2a, Extended Data Fig. 3a**). Among the differentially expressed genes, 3-hydroxymethylglutaryl-CoA synthase 2 (*Hmgcs2*) – the rate-limiting enzyme in ketogenesis – was markedly upregulated in ISCs from KD-fed mice (**Fig. 2b, c**). This finding is consistent with prior observations that *Hmgcs2* expression is enriched within Lgr5+ ISCs, with expression declining as cells exit the stem cell compartment and differentiate, and that loss of *Hmgcs2* impairs ISC differentiation and regenerative capacity, effects that can be partially rescued by KD exposure or exogenous beta-hydroxybutyrate (BHB) administration ^18^.

**Figure 2.**
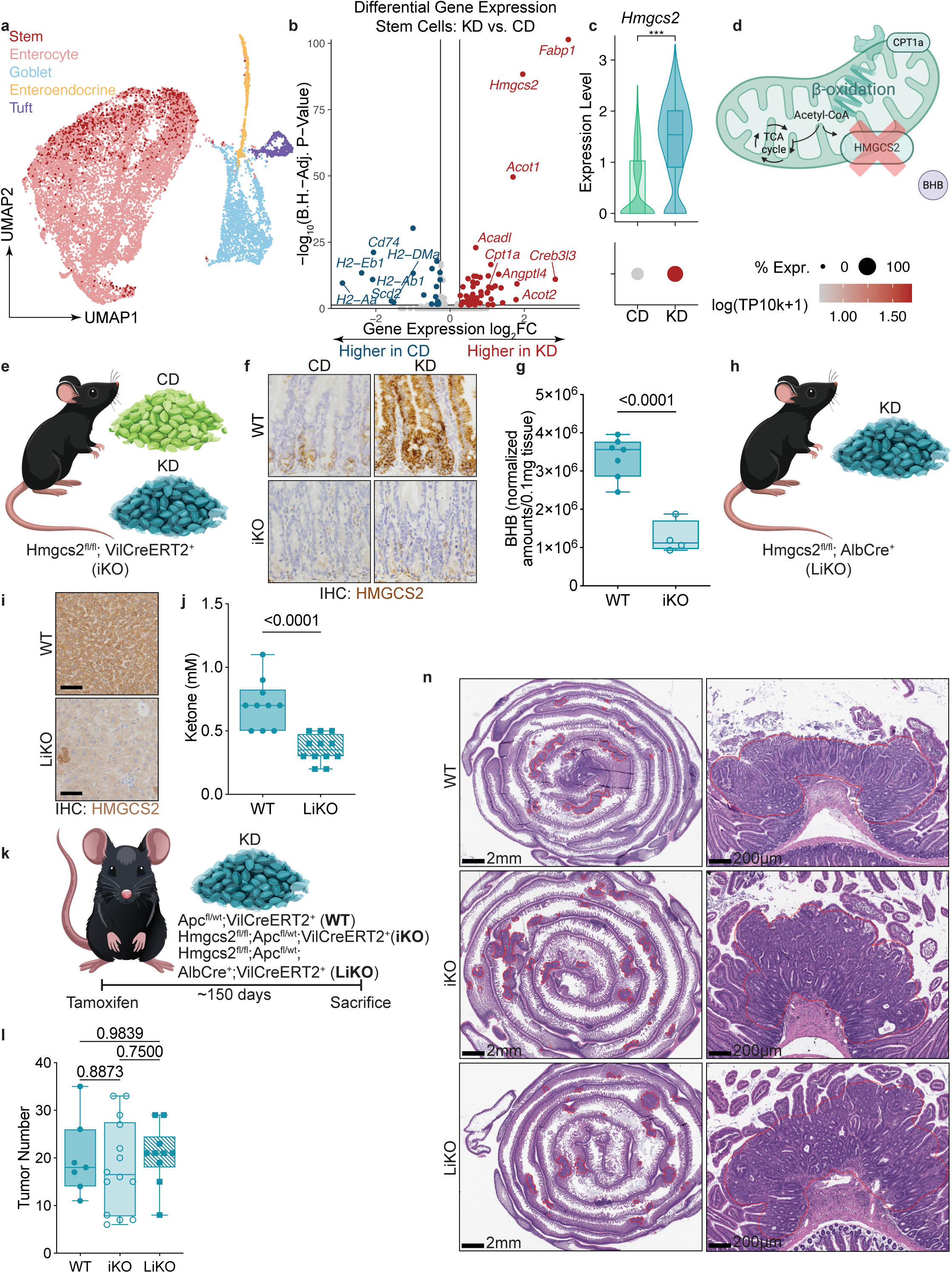
Loss of epithelial or systemic ketogenesis does not mitigate the pro-tumorigenic effects of a ketogenic diet. (a) UMAP visualization of 15,053 single cells isolated from the intestinal epithelium of control diet (CD) and ketogenic diet (KD) mice, colored by cell type. (b) Differential gene expression of ISCs from KD- and CD-fed mice (c) Hmgcs2 expression in ISCs from CD and KD mice. (d,e) Schematic of beta-oxidation and ketogenesis with inducible Hmgcs2 knockout (iKO) models (f) Confirmation of Hmgcs2 deletion in the intestinal epithelium of iKO mice by immunohistochemistry. (g) Quantification of beta-hydroxybutyrate (BHB) levels in intestinal tissue of KD-fed WT and iKO mice. (h) Schematic of hepatocyte-specific Hmgcs2 knockout (LiKO) mice. (i) Immunohistochemistry for HMGCS2 demonstrating hepatocyte-specific loss of HMGCS2 expression in LiKO mice. (j) Serum ketone levels under KD conditions in LiKO and control mice. k) Breeding scheme to generate WT, iKO, and LiKO mice for tumorigenesis studies. (l) Quantification of tumour number and tumour burden across CD and KD conditions in wild-type, iKO, and LiKO mice with (m) representative H&E images. Data are mean ± s.d.

To directly investigate the contribution of epithelial-derived ketone bodies, we generated an inducible intestinal-specific Hmgcs2 knockout model (iKO) by crossing Vil-CreERT2 mice with Hmgcs2^fl/fl^ mice, enabling deletion of Hmgcs2 in all intestinal epithelial cells following tamoxifen administration (**Fig. 2d, e**). KD-induced upregulation of epithelial HMGCS2 was abolished in iKO mice, confirming efficient deletion (**Fig. 2f**). Loss of epithelial HMGCS2 significantly reduced intestinal BHB production without appreciably altering local lipid species, indicating that a substantial fraction of ketone bodies within the intestinal epithelium are produced locally rather than being derived from the circulation and that lipid flux was unlikely to be significantly impacted (**Fig. 2g, Extended Data Fig. 2b**). To complement these findings and examine systemic contributions, we utilized Alb-Cre mice, which constitutively express Cre recombinase under the Albumin promoter, to delete Hmgcs2 specifically in hepatocytes, generating liver-specific knockout (LiKO) mice (**Fig. 2h, i, Extended Data Fig. 3c–f**). LiKO mice exhibited dramatically decreased circulating ketone levels under both fasting and ketogenic dietary conditions (**Fig. 2j**), thereby enabling us to model reductions in systemic ketogenesis.

To specifically evaluate the role of local and systemic ketogenesis on tumour development with KD feeding, we crossed both Hmgcs2 iKO and LiKO models into the established Apc^fl/wt^ heterozygous background, which sensitizes mice to intestinal adenoma formation following single allele loss of Apc (**Fig. 2k**). As expected, the liver exhibited no significant histopathologic changes following loss of a single Apc allele, likely due to differences in proliferative kinetics between hepatic and intestinal tissues. Based on prior work suggesting a tumor-suppressive effect of BHB, we hypothesized that loss of ketogenesis would exacerbate KD-induced tumour burden. Surprisingly, however, neither intestinal (iKO) nor systemic (LiKO) loss of ketone body production mitigated the pro-tumorigenic effects of a ketogenic diet (**Fig. 2l-m**).

Given that ketone body metabolites can also serve as energetic substrates via ketolysis (i.e. a process whereby ketone bodies are converted to acetyl-CoA), we next investigated whether impairing ketone body utilization would affect tumour burden. BHB is oxidized to acetoacetate (AcAc) by the mitochondrial enzyme 3-hydroxybutyrate dehydrogenase 1 (BDH1) and further activated to acetoacetyl-CoA by the enzyme succinyl-CoA:3-ketoacid CoA transferase 1 (OXCT1), enabling entry into the tricarboxylic acid (TCA) cycle for energy production. Unlike HMGCS2, which displays a restricted expression profile, both BDH1 and OXCT1 were found to be broadly expressed throughout the intestinal epithelium (**Extended Data Fig. 4a-b**). Loss of BDH1 and OXCT1 allows manipulation of the ketogenic/ketolytic pathways without altering KD-induced HMGCS2 expression (**Extended Data Fig. 4c**). To disrupt ketolytic capacity, we crossed Bdh1^fl/fl^ and Oxct1^fl/fl^ mice into the Apc^fl/wt^ background (**Extended Data Fig. 4d**).

Consistent with observations from the HMGCS2 iKO models, loss of BDH1 or OXCT1 did not significantly impact tumour burden (**Extended Data Fig. 4d-f**). These data further demonstrate that neither canonical ketogenesis nor ketolytic pathways in epithelial cells or systemically are required to promote intestinal tumorigenesis in the context of a ketogenic diet. Together, these findings highlight that the pro-tumorigenic effects associated with KD feeding are likely mediated through lipid-driven mechanisms independent of local or systemic ketone body metabolism.

### Induction of ectopic ketogenesis is insufficient to modulate intestinal tumorigenesis

A ketogenic diet is characterized by low carbohydrate content and is primarily defined by its high fat composition. To isolate the effects of increased ketone body production from the confounding influence of high dietary lipid intake, we generated a novel Rosa26-LSL-Hmgcs2-2A-mCherry mouse model (hereafter referred to as HMGCS2 iOE), wherein Hmgcs2 expression is placed under Cre recombinase control (**Fig. 3a, Extended Data Fig. 5a**). By crossing these mice to the tamoxifen-inducible Vil-CreERT2 line, we achieved spatial and temporal control of Hmgcs2 expression specifically within the intestinal epithelium, thereby allowing modulation of ketone body production independent of dietary manipulation. Upon tamoxifen induction, HMGCS2 iOE mice exhibited robust and widespread expression of both HMGCS2 protein and the linked mCherry reporter in intestinal epithelial cells (**Fig. 3b, Extended Data Fig. 5b**).

**Figure 3.**
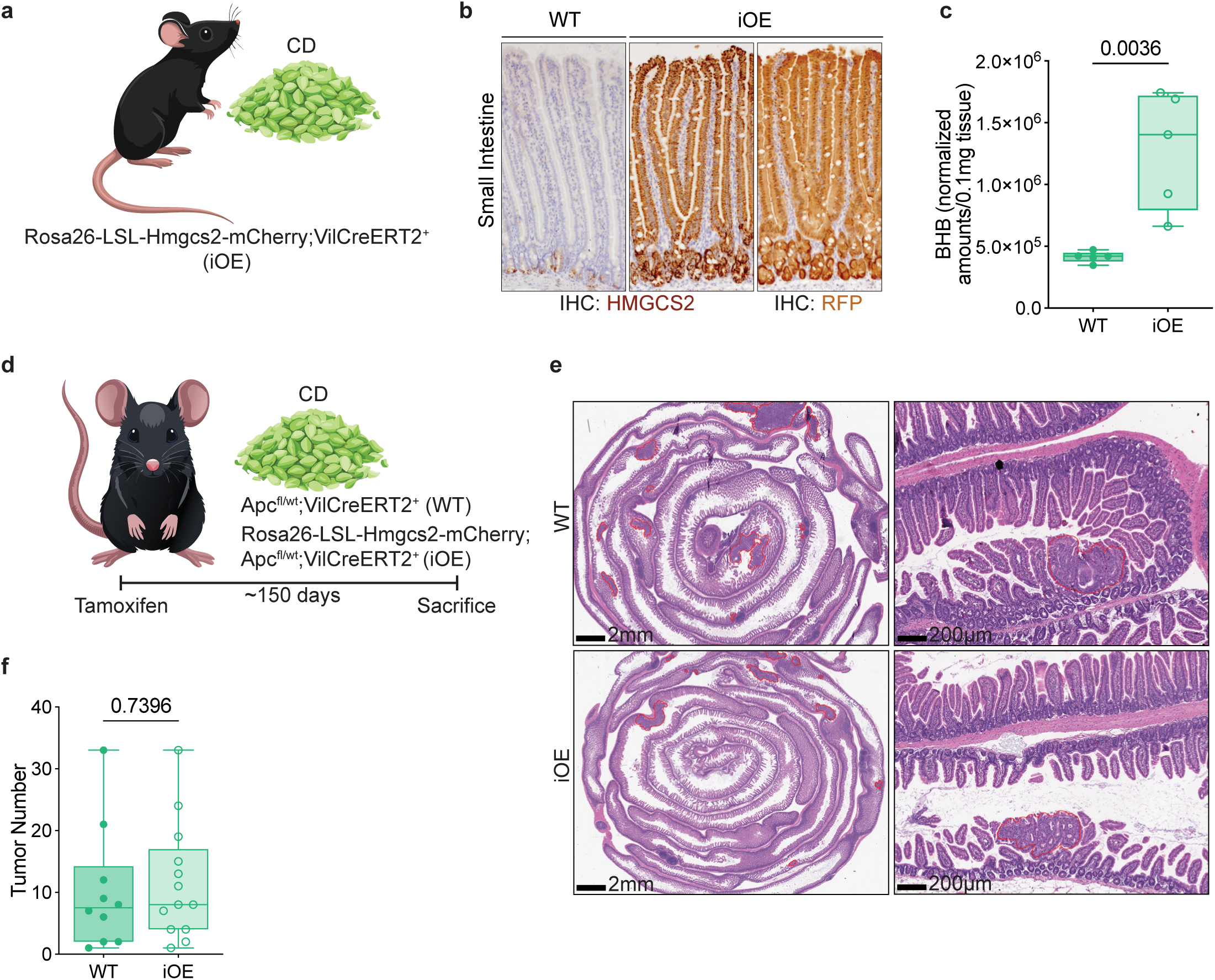
Tissue-specific Hmgcs2 overexpression elevates intestinal ketone body levels without affecting spontaneous tumorigenesis. (a) Schematic of the Rosa26-CAG-Hmgcs2-2A-mCherry-LSL (HMGCS2 iOE) mouse (b) Representative immunofluorescence images showing expression of HMGCS2 and mCherry throughout the intestinal epithelium following Vil-CreERT2-mediated induction. (c) Quantification of β-hydroxybutyrate (BHB) levels in intestinal tissue lysates from control diet-fed HMGCS2 iOE mice and WT controls. (d) Breeding scheme to generate mice for tumorigenesis studies with or without concurrent Hmgcs2 overexpression maintained on control diet. (e) Tumour burden quantification for WT and iOE mice. (f) Representative images of tumour burden along the intestinal tract. Data are mean ± s.d.

Importantly, ectopic overexpression of Hmgcs2 in the intestinal epithelium led to a significant elevation of local BHB concentrations compared to control animals, despite maintenance on a standard control diet (**Fig. 3c, Extended Data Fig**. **5c**). These findings confirm that tissue-specific overexpression of HMGCS2 is sufficient to drive augmented ketone body production *in vivo* without the need for a lipid-enriched or carbohydrate-restricted diet. Thus, the HMGCS2 iOE model enables dissection of the epithelial-intrinsic effects of increased ketone metabolism in the absence of systemic metabolic alterations associated with ketogenic or high-fat diets.

To determine whether enhanced epithelial ketone body production influences tumorigenesis, we next assessed spontaneous adenoma formation in Vil-CreERT2; Apc^fl/wt^ mice with or without concurrent Hmgcs2 overexpression, all maintained on a purified control diet. Surprisingly, enforced Hmgcs2 expression, which increased intestinal ketone levels (**Fig. 3c**) did not significantly alter tumour numbers, tumour sizes, or survival compared to wild-type littermates (**Fig. 3d-f**). These data indicate that while epithelial ketone body production can be augmented independently of dietary fat intake, such augmentation alone is insufficient to modify basal intestinal tumorigenesis or clonogenic potential under homeostatic conditions.

### PPAR-mediated long-chain FAO contributes to KD-associated intestinal tumorigenesis

To define the impact of a KD on ISC molecular states and metabolism, we conducted transcriptomic analyses of ISCs from mice maintained on either a KD or a control diet (CD). Using CollecTRI, we aimed to identify transcription factors whose genome-wide targets changed in expression with KD. KD exposure significantly increased activation of multiple peroxisome proliferator-activated receptors (PPARs) and PPAR gamma coactivator 1-beta (Ppargc1b) (**Fig. 4a**). Additionally, we found that SMADs and SREBF2 were predicted to have increased activity. In contrast, RFXAP/RFANK were predicted to have less activity, suggesting a potential tradeoff with the upregulation of programs involved in lipid metabolism and stem cell maintenance at the expense of MHC II expression. Gene set enrichment analysis (GSEA) revealed robust enrichment of lipid catabolism and fatty acid oxidation (FAO) pathways, including a marked induction of genes involved in mitochondrial long-chain fatty acid transport and β-oxidation with corresponding downregulation in antigen presentation (**Fig. 4b, Extended Data 6a**). We complemented these findings by specifically scoring for markers of PPARs from multiple sources (**Fig. 4c-e**). Among FAO-associated transcripts, carnitine palmitoyltransferase 1A (*Cpt1a*), which catalyzes the rate-limiting step in mitochondrial import of long-chain fatty acids for downstream oxidation, was significantly elevated in KD ISCs compared to CD ISCs (**Fig. 4f**). Immunoblotting confirmed increased CPT1a protein expression in crypt epithelial isolates following KD feeding (**Extended Data Fig. 1i**).

**Figure 4.**
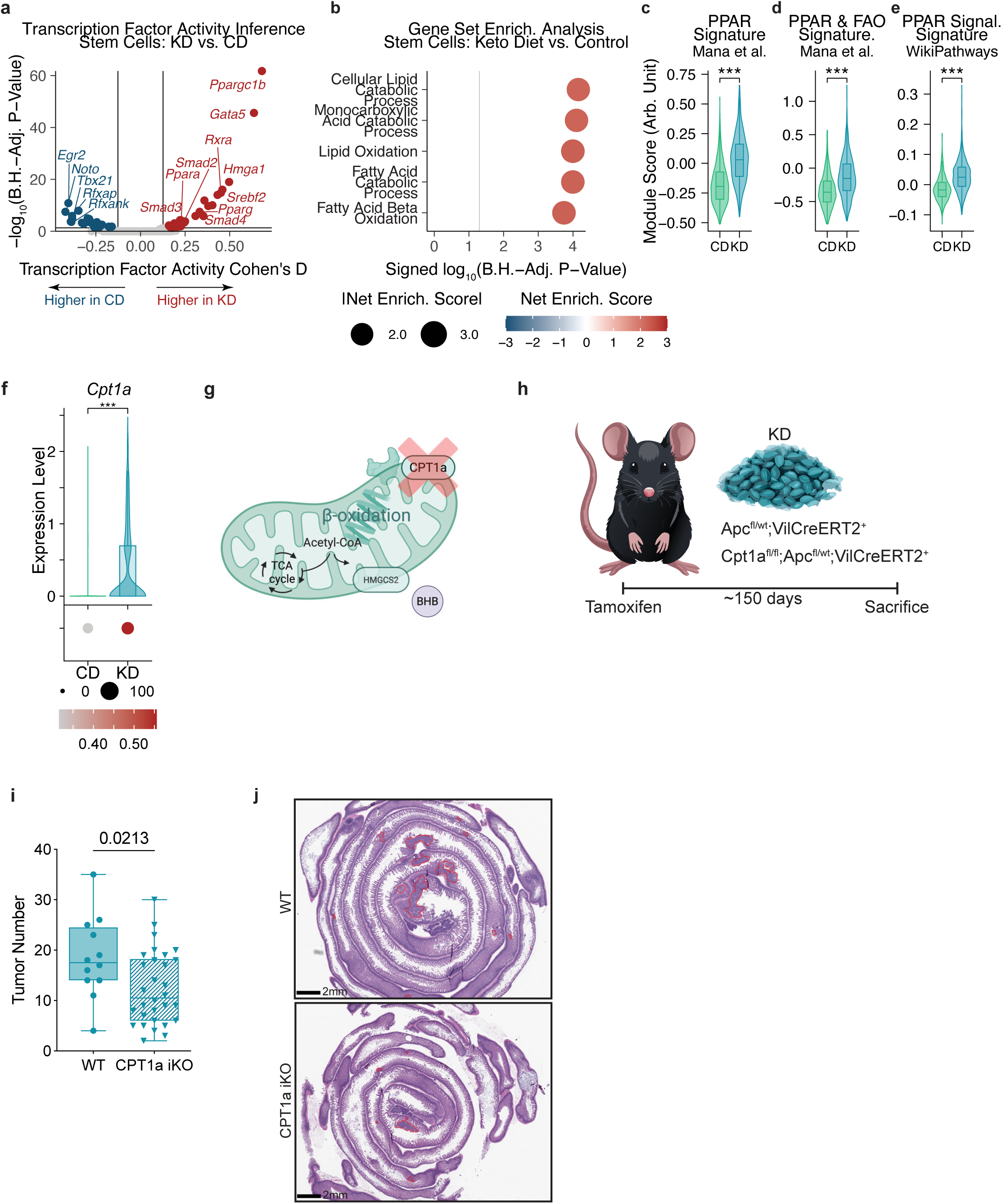
Ketogenic diet enhances PPAR signaling, fatty acid oxidation gene expression, and ISC function via CPT1a activity. (a) Transcription factor activity inference for ISCs in CD- and KD-fed mice (b) Gene set enrichment analysis (GSEA) of most increased pathways in ISCs isolated from KD- and CD-fed mice (c-e) PPAR signature scoring for CD- and KD-ISCs (f) Cpt1a expression in CD- and KD-ISCs (g,h) Schematic of beta-oxidation and ketogenesis with inducible CPT1a knockout models (i) Representative H&E images with (j) quantification of total tumour number in WT and CPT1a iKO mice. Data are mean ± s.d.

To determine the functional relevance of FAO in KD-associated tumorigenesis, we crossed Cpt1a^fl/fl^ mice to Vil-CreERT2;Apc^fl/wt^ mice to generate inducible intestinal-specific Cpt1a knockout (Cpt1a iKO) animals (**Fig. 4g,h)**. Following tamoxifen-induced recombination, cohorts were maintained on KD or CD diets, and spontaneous tumour formation was assessed over time. In the Apc-het background, Cpt1a deletion significantly reduced tumour burden in KD-fed mice compared to WT controls. Quantitative analyses revealed a reduction in the total number of tumours in the Cpt1a iKO cohort in the presence of KD but not CD (**Fig. 4i-j, Extended Data Fig. 6b-e**). Together, these findings indicate that ketogenic diet-induced PPAR activation enhances ISC function through an FAO-dependent mechanism, with CPT1a acting as a critical metabolic effector. Loss of CPT1a disrupts this pathway, attenuating the KD-associated increase in tumorigenesis. Of note, similar findings are seen when Cpt1a^fl/fl^; Apc^fl/wt^; Lgr5-EGFP-IRES-CreERT2 mice are exposed to a diet with high lipid content ^13^. These data highlight mitochondrial FAO rather than ketogenesis or ketone body metabolites as a key node linking dietary lipid exposure, ISC metabolic adaptation, and intestinal cancer risk.

## Discussion

Here, we demonstrate that a ketogenic diet (KD), despite its physiologic and hormonal profile as distinct from an obesogenic high-fat diet (HFD), shares key tumorigenic features. Through a series of genetic and dietary manipulations, we dissect the contributions of ketogenesis and ketone body metabolism to intestinal tumorigenesis, revealing that dietary lipid and resulting PPAR-mediated FAO, rather than ketone body production, is the principal influencer for KD effects on intestinal tumorigenesis. These findings indicate that while ketone body production is modifiable through diet and genetic perturbation, ketone body metabolites are neither necessary nor sufficient to mediate KD-induced tumorigenesis.

In the mouse colon, HMGCS2 expression and ketogenesis are additionally influenced by the resident microbiome (**Extended Data 7a**) ^40^. When compared to a purified control diet however, a ketogenic diet increases epithelial beta-hydroxybutyrate (**Extended Data 7b**). This can also be modulated through overexpression of HMGCS2 as with the iOE mouse (**Extended Data 7c-d**). The dietary influence of KD on colonic tumour burden in this model is the opposite of the small intestine and yet remains independent of ketone bodies: a KD results in fewer colonic tumours and genetic modulation of epithelial ketones has no discernible anti-tumorigenic effects (**Extended Data 7e-f**). Again, the G-protein coupled receptor HCAR2 has been previously postulated as a key intermediary however, expression appears largely restricted to normal epithelium and is absent from both CD and KD colonic tumours (**Extended Data 7g**). Although beyond the scope of this work, there are numerous changes that occur in the colonic milieu when exposed to high lipid content and these could, in combination, explain some of the observed effects. This work challenges the prevailing notion that ketone bodies are uniformly beneficial or protective against cancer, revealing that in the intestinal epithelium, their effects are diet-modulated yet context-dependent.

Importantly, our observations reconcile previous paradoxes in the field. Dietary interventions, such as fasting, caloric restriction, and exogenous ketone supplementation have been associated with enhanced ISC regenerative capacity and reduced tumorigenesis. However, they differ fundamentally from KD in that they lack sustained lipid overload. In a similar manner, the post-fast refeeding exists as a distinct state that can also lead to increased tumour incidence ^24^ . Thus, the diet composition and systemic metabolic effects must be considered when comparing opposing influences on tumorigenesis such as chronic lipid-induced FAO engagement seen with KD. In conclusion, our findings reveal that a high-fat ketogenic diet influences intestinal tumorigenesis through lipid-driven metabolic reprogramming rather than ketone body signaling, stimulating FAO activation at the earliest stages of tumour initiation. These insights have important implications for dietary interventions in cancer prevention and treatment, particularly in individuals with hereditary cancer predisposition syndromes such as FAP who are prone to small intestinal tumorigenesis, and highlight the complex interplay between metabolism and oncogenesis in the intestinal epithelium.

## Author Contributions

J.E.S.S. and F.C. initiated the project, conceived, designed, performed, and interpreted all experiments and wrote the manuscript with help from O.H.Y.. C.T., J.T.H.S., K.J.W., T.S., and I.F. performed experiments, assisted with data collection and interpretation. J.T.H.S. assisted with intestinal tissue metabolomic analysis. K.J.W. assisted with intestinal tissue lipidomic analysis. All authors assisted in interpreting experiments, writing, and editing the manuscript.

## Ethics Declaration

Competing interests: The authors declare no competing interests.

## Acknowledgements

We thank the Swanson Biotechnology Center at the Koch Institute, including the Flow Cytometry, Histology, and ES Cell and Transgenics core facilities; the Department of Comparative Medicine for mouse husbandry support; S. Holder and members of the Hope Babette Tang (1983) Histology Facility for substantial histological support; all the members of the Yilmaz laboratory for discussions; K. Kelley for laboratory management; and L. Galoyan for administrative assistance. Ö.H.Y. is supported by the National Institutes of Health (5R01CA245314, R01CA211184, R01CA034992, R01CA257523, R01DK126545, U01CA250554 and U54CA224068); a Pew-Stewart Trust scholar award; the Kathy and Curt Marble cancer research award; a Koch Institute– Dana-Farber/Harvard Cancer Center Bridge project grant; and AFAR. Ö.H.Y. receives support from the MIT Stem Cell Initiative. J.E.S. is supported by an American Gastroenterological Association Research Scholar Award (AGA2025-13-05). F.C. is supported by a Damon Runyon Postdoctoral Fellowship Award (DRG-2463-22).

## Methods

### Mouse Strains

Mice were under the husbandry care of the Department of Comparative Medicine at the Koch Institute for Integrative Cancer Research at Massachusetts Institute of Technology. All procedures were conducted according to AALAC guidelines and approved by the MIT Committee on Animal Care. Lgr5-2A-EGFP-2A-CreERT2 were created and previously described ^39^. Lgr5-eGFP-IRES-CreERT2 (strain name: B6.129P2-Lgr5^tm1(cre/ERT2)Cle^/J; stock number 008875) mice were purchased from Jackson Laboratory. Apc-het mice were created by crossing the previously described Apc*^loxp^* exon 14 (Apc*^fl/fl^*) mice to mice carrying a wild type Apc allele ^41^. Villin-CreERT2 (Vil-CreERT2) mice were a gift from S. Robine and previously described ^42^. Hmgcs2*^fl/fl^* were originally generated and described previously ^18^. Albumin-Cre mice (strain name B6.Cg-*Speer6-ps1^Tg(Alb-cre)21Mgn^*/J; stock number #003574) were purchased from Jackson Laboratory. Bdh1*^loxp/loxp^* mice were a gift from Daniel Kelly and described previously ^43^. Oxct1*^fl/fl^* mice were a gift from Peter Crawford and previously described ^44^. Rosa26-LSL-Hmgcs2-mCherry mice were created by co-injecting Cas9 mRNA and a donor vector containing “CAG promoter-loxP-3*polyA-loxP-Kozak-mHmgcs2 CDS-2A-mCherry-polyA” cassette into fertilized eggs to generate targeted conditional knockin offspring. Cpt1a *^fl/fl^* mice were a gift from Peter Carmeliet and described previously ^45^. The following strains were bred in-house: 1) Lgr5-2A-EGFP-2A-CreERT2 , 2) Lgr5-eGFP-IRES-CreERT2, 3) Apc^fl/wt^; Villin-CreERT2, 4) Hmgcs2^fl/fl^; Villin-CreERT2, 5) Hmgcs2^fl/fl^; Albumin-Cre, 6) Hmgcs2^fl/fl^; Apc^fl/wt^; Villin-CreERT2, 7) Hmgcs2^fl/fl^; Apc^fl/wt^; Villin-CreERT2; Albumin-Cre, 8) Bdh1^fl/fl^; Apc^fl/wt^; Villin-CreERT2, 9) Oxct1^fl/fl^; Apc^fl/wt^; Villin-CreERT2, 10) Rosa26-LSL-Hmgcs2-mCherry; Villin-CreERT2, 11) Rosa26-LSL-Hmgcs2-mCherry; Apc^fl/wt^; Villin-CreERT2, 12) CPT1a^fl/fl^; Apc^fl/wt^; Villin-CreERT2. All mice were age and sex-matched with both male and female mice used. Food was provided *ad libitum*. Control diet with 10% fat (Research Diet D12450J) and ketogenic diet with 80% fat (Research Diet D06040601) were used throughout experimental conditions with the exception of metabolomic profiling of conventional (CONV) and Hmgcs2 iOE mice where standard chow (LabDiet, 5P76) was used.

### In vivo treatments

Tamoxifen-induced Cre activation was obtained through intraperitoneal tamoxifen injection suspended in 1 part 100% ethanol to 9 parts sunflower seed oil (Spectrum S1929) at a concentration of 10mg/mL and dosed at 100mg/kg. For tumour studies, mice harboring conditional alleles plus Apc*^fl/wt^*were administered tamoxifen two times within one week. For additional non-tumor studies, mice were administered tamoxifen three times within one week. BrdU (sigma Aldrich, 19-160) was prepared at 10mg/mL in PBS and injected at 100mg/kg 4 hours prior to tissue collection.

### Tumor analysis

Intestine was dissected out and flushed with 4°C PBS before being placed in histology cassettes and fixed overnight in 10% formalin. Tissue was then transferred to 70% ethanol before additional dehydration steps and embedding. 4-5μm sections were stained with hematoxylin and eosin and scanned in at 20x magnification using Leica Aperio Slide Scanner. SVS files were then analyzed using QuPath in a blinded manner.

### Serum measurements

Circulating glucose and ketones were measured by collecting tail vein blood and using FreeStyle Precision Neo System Kit with Blood Glucose and Ketone Test Strips (Abbott). Plasma was collected via cardiac puncture into capillary blood collection vial (Microvette 20.1292.100) before separation via centrifugation. Circulating insulin was then quantified via wide-range ultra sensitive ELISA (Crystal Chem #90080).

### Tissue metabolic profiling

For metabolomics, intestine tissue was quickly dissected out, washed twice with 150mM Ammonium acetate, and snap-frozen into liquid nitrogen. The snap-frozen tissue was then homogenized into powder using mortar-pestle in liquid nitrogen. About 50mg of the tissue powder was used for metabolites extraction in 80% MeOH with OMNI bead homogenizer. The tissue/80%MeOH mixture was incubated for 24 hours at −80°C and then centrifuged (16,000 g) for 20 minutes at 4°C. The MeOH supernatants were transferred into new tubes and placed on ice. The sample pellets were resuspended in 500μl 0.2M NaOH, heated at 95°C for 10 minutes, and the protein concentration was measured by BCA assay using 1:10 dilution. Finally, the 500 μg protein equivalent of MeOH supernatants were dried using multivap nitrogen evaporator. Dried metabolites were resuspended in 50% ACN:water at 50 mg tissue extract/ml. 5μl was loaded onto a Luna NH2 3um 100A (150 × 2.0 mm) column (Phenomenex) using a Vanquish Flex UPLC (Thermo Scientific). The chromatographic separation was performed with mobile phases A (5 mM NH4AcO pH 9.9) and B (ACN) at a flow rate of 200 μl/min. A linear gradient from 15% A to 95% A over 18 min was followed by 7 min isocratic flow at 95% A and reequilibration to 15% A. Metabolites were detected with a Thermo Scientific Q Exactive mass spectrometer run with polarity switching in full scan mode using a range of 70-975 m/z and 70.000 resolution. Maven (v 8.1.27.11) was used to quantify the targeted polar metabolites by AreaTop, using expected retention time and accurate mass measurements (< 5 ppm) for identification. Data analysis, including principal component analysis and heat map generation was performed using in-house R scripts.

For lipidomics, isolated intestine tissue was snap frozen in liquid nitrogen. 50mg of homogenized tissue was then sent to UCLA Lipidomics in 2mL homogenizer tubes pre-loaded with (6) 2.8mm ceramic beads (Omni #19-628). 750ul PBS was added to the tube and homogenized in the Omni Bead Ruptor Elite (3 cycles of 10 seconds at 5 m/s with a 10 second dwell time). Homogenate containing 1-8mg of original tissue was transferred to a glass tube for extraction. A modified Bligh and Dyer extraction was carried out on all samples ^46^ . Prior to biphasic extraction, a standard mixture of 75 lipid standards based on Avanti Ultimate Splash ONE mix was used (Avanti 330820, 861809, 330729, 330727, 791642, 330726). Following two successive extractions, pooled organic layers were dried down in a Thermo SpeedVac SPD300DDA using ramp setting 4 at 35 degrees C for 45 minutes with a total run time of 90 minutes. Lipid samples were resuspended in 1:1 methanol/dichloromethane with 10mM Ammonium Acetate and transferred to robovials (Thermo 10800107) for analysis. Samples were analyzed on the Sciex 6500+ with DMS device with an expanded targeted acquisition list consisting of 1450 lipid species across 17 subclasses. Differential Mobility Device was tuned with EquiSPLASH LIPIDOMIX (Avanti 330731). Data analysis performed on an in-house data analysis platform comparable to the Lipidyzer Workflow Manager with instrument method including settings, tuning protocol, and MRM list as described ^47^. Quantitative values were normalized to mg of tissue.

### Crypt and ISC isolation and culturing

Intestine was removed and flushed with 4°C PBS (no calcium, no magnesium), opened laterally and gently cleaned and cut into sections before rinsing and incubation at 4°C in 10mM EDTA for 40 minutes. Crypts were then mechanically separated by shaking and filtering through 70um mesh to remove villi and tissue fragments. Isolated crypts and flow isolated ISCs were counted and embedded in Matrigel (Corning 356231) as previously described ^13^.

### Flow cytometry

Following crypt isolation, crypt suspensions were pelleted (100g, 5 min, 4°C) and supernatant was discarded. Crypts were resuspended in TrypLE (GIBCO, no phenol red, 1264039) and dissociated into individual cells by warming crypts in a 32°C water bath for 60 seconds. Dissociated single cells were incubated with the following antibodies for flow cytometry analysis: CD45-PE (eBioscience, 12-0451-83), CD31-PE (BioLegend, 102514), Ter-119PE (Biolegend, 116208), CD24-Pacific Blue (Biolegend, 101820), CD117-APC/Cy7 (Biolegend, 105826), and EpCAM-APC (eBioscience, 17-5791-82). 7AAD (Invitrogen, A1310) was used as a viability dye to exclude dead cells from analysis. Intestinal stem cells (ISCs) and progenitors were isolated as Lrg5-eGFP^hi^EpCAM^+^CD24^low/-^CD31^-^Ter119^-^CD45^-^7AAD^-^ and Lrg5-eGFP^low^EpCAM^+^CD24^low/-^CD31^-^Ter119^-^CD45^-^7AAD^-^, respectively. Cells were sorted using a BD FACS II SORP cell sorter.

### scRNA-seq processing

Cells were sorted with the same parameters as described above for flow-cytometry into an Eppendorf tube containing 50ml of 0.4% BSA-PBS and stored on ice until proceeding to the Chromium Single Cell Platform. Single cells were processed through the Chromium Single Cell Platform using the Chromium Single Cell 3’ Library, Gel Bead and Chip Kits (10X Genomics, Pleasanton, CA), following the manufacturer’s protocol. Briefly, an input of 7,000 cells was added to each channel of a chip with a recovery rate of 1,500-2,500 cells. The cells were then partitioned into Gel Beads in Emulsion (GEMs) in the Chromium instrument, where cell lysis and barcoded reverse transcription of RNA occurred, followed by amplification, tagmentation and 50 adaptor attachment. Libraries were sequenced on an Illumina NextSeq 500.

The CellRanger pipeline was used to align sequencing reads to the mm10 genome according to manufacturer recommendations. Cells were pre-processed to remove cells with fewer than 1,750 UMIs detected (likely corresponding to detection of ambient RNA or low-quality cells) or more than 75,000 UMIs (likely corresponding to doublets). Data was then analyzed using Seurat in R with normalization implemented via SCTransform’s regularized negative binomial regression. For downstream visualization and clustering, principal component analysis was performed on the highly variable genes returned from SCTransform, with the number of principal components made by examining the elbow plot of variance explained by each principal component (results robust against alternate choices of QC and visualization parameters). UMAP visualization was implemented using the ‘‘uwot’’ R package. Stem cells were identified using a module score of core intestinal stem cell genes calculated with Seurat’s ‘‘AddModuleScore’’ function (*Lgr5*, *Axin2*, *Olfm4*, *Lrig1*, *Fzd7*), as previously described ^13^. Stem cells were additionally scored for modules of PPAR target genes (*Hmgcs2*, *Angptl4*, *Pdk4*, *Acsl3*, *Me1*, *Acot1*, *Reg3g*, *Plin2*), fatty acid oxidation-related genes regulated by PPAR (*Cpt1a*, *Acadvl*, *Cact*, *Acadl*, *Hadha*, *Hadhb*, *Eci2*), and the WikiPathways PPAR Signaling gene set, as previously described ^13^. GSEA was run through the fgsea package ^48^ . CollecTRI was run through the decoupleR package, and effect sizes were calculated as signed Cohen’s D effect sizes ^49^. Statistical significance was calculated for differential gene expression, differential module scores, and differential transcription factor activities using the non-parametric Mann-Whitney test and adjusted for multiple hypothesis testing using the Benjamini-Hochberg correction.

### Immunohistochemistry and Immunofluorescence

Tissue was fixed in 10% normal buffered formalin prior to paraffin-embedding. 4-5μm micron sections underwent deparaffinization and rehydration prior to antigen retrieval using Borg Decloacker RTU solution (Biocare Medical, BD1000G1) and a pressurized Decloaking Chamber (Biocare Medical, NxGen). Antibodies and respective dilutions used for immunohistochemistry are as follows: goat polyclonal anti-GFP (1:1000, Abcam, ab6673), rat anti-BrdU (1:2000, Abcam, 6326), rabbit polyclonal anti-lysozyme (1:2000, ThermoFisher, RB-372-A1), rabbit monoclonal anti-HMGCS2 (1:2000, Abcam, ab137043), rabbit polyclonal anti-OXCT1 (1:1000, Sigma-Aldrich, HPA012047), rabbit polyclonal anti-BDH1 (1:500, Proteintech, 15417-1-AP), with dilutions performed in Signalstain Antibody Diluent (Cell Signaling Technology, #8112). Biotin conjugated secondary donkey anti-rabbit antibodies (1:500, Jackson ImmunoResearch) were used prior to Vectastain Elite ABC immunoperoxidase detection kit (Vector Laboratories, PK6100). Visualization was performed using Signalstain DAB substrate kit (Cell Signaling Technology, #8049). Counterstaining was performed with 50% Gill No. 1 hematoxylin solution (Sigma-Aldrich, GHS116). Images were acquired using an Olympus BX43 microscope with 4x and 10x objectives (Olympus UPlanSApo). The following primary antibodies were used for immunofluorescence: rabbit polyclonal anti-HCAR2 (1:200, ThermoFisher, PA5-50671) and mouse monoclonal anti-beta-catenin (1:200, BD Biosciences, 610164). Alexa Fluor secondary antibodies, anti-mouse 488 and anti-rabbit 568 were use for visualization. Tissue was mounted using Invitrogen Prolong Gold mounting medium containing DAPI. Images were acquired using an Olympus FV1200 Laser Scanning Confocal Microscope.

### Immunoblotting

Snap frozen tissue was crushed into a fine powder before elution with RIPA buffer and tissue homogenizer. Isolated crypts were resuspended in RIPA buffer for lysis. In both cases, after centrifuging at 10,000 rpm for 15 minutes at 4°C, the resulting supernatant was quantified using the Pierce BCA protein assay kit (23225, Thermo Fisher Scientific) then suspended in Laemmli SDS sample buffer (J61337-AC, VWR) at a concentration of 2ug/ul and boiled for 5 minutes at 95°C. 20ug protein per sample was separated on 4-12% gradient gel (NuPAGE Bis-Tris Mini Protein, Invitrogen, NP0336BOX) and transferred onto a 0.45 micron PVDF membrane (Immobilon-P transfer, Millipore, ipvh00010) and analyzed by immunoblotting using rabbit monoclonal anti-HMGCS2 (1:1000, Abcam, ab137043), mouse monoclonal anti-CPT1a (1:1000, Abcam, ab128568), rabbit polyclonal anti-BDH1 (1:1000, Proteintech, 15417-1-AP), rabbit polyclonal anti-OXCT1 (1:1000, Sigma-Aldrich, HPA012047), rabbit monoclonal anti-vinculin (1:1000, Cell Signaling Technology, #13901) followed by horseradish peroxidase (HRP)-conjugated IgG secondary antibodies (1:3000, Cell Signaling Technology, #7076, #7074) and ECL detection kit (ECL K-12045-D20 and Sirius K-12043-D20, Advansta). Quantitative chemiluminescent imaging was performed using the ImageQuant LAS 4000 (GE Healthcare).

### Statistics and Reproducibility

Unless otherwise specified in the main text or figure legends, all experiments reported in this study were repeated at least three independent times. Unless otherwise specified in the main text or figure legend, all sample numbers (n) represent biological replicates. No sample or animals were excluded from analysis, and sample size estimates were not used. Studies were not conducted blind with the exception of all histological analyses. Age-matched and sex-matched mice were randomly assigned to groups. All values are presented as mean ± s.d. Unless otherwise specified in the figure legend, intergroup comparison was performed using unpaired two-tailed t-tests. In the case of more than two comparison groups, ANOVA with Tukey’s was performed. Statistical analysis was performed using GraphPad Prism 10. Statistical details can be found in the figure legends.

### Data availability

Datasets generated in this study will be made publicly available upon publication.

