## Supplementary figures and images for "Dietary lipids, not ketone body metabolites, influence intestinal tumorigenesis in a ketogenic diet"

### Extended Data Figure 1

#### Extended Data 1

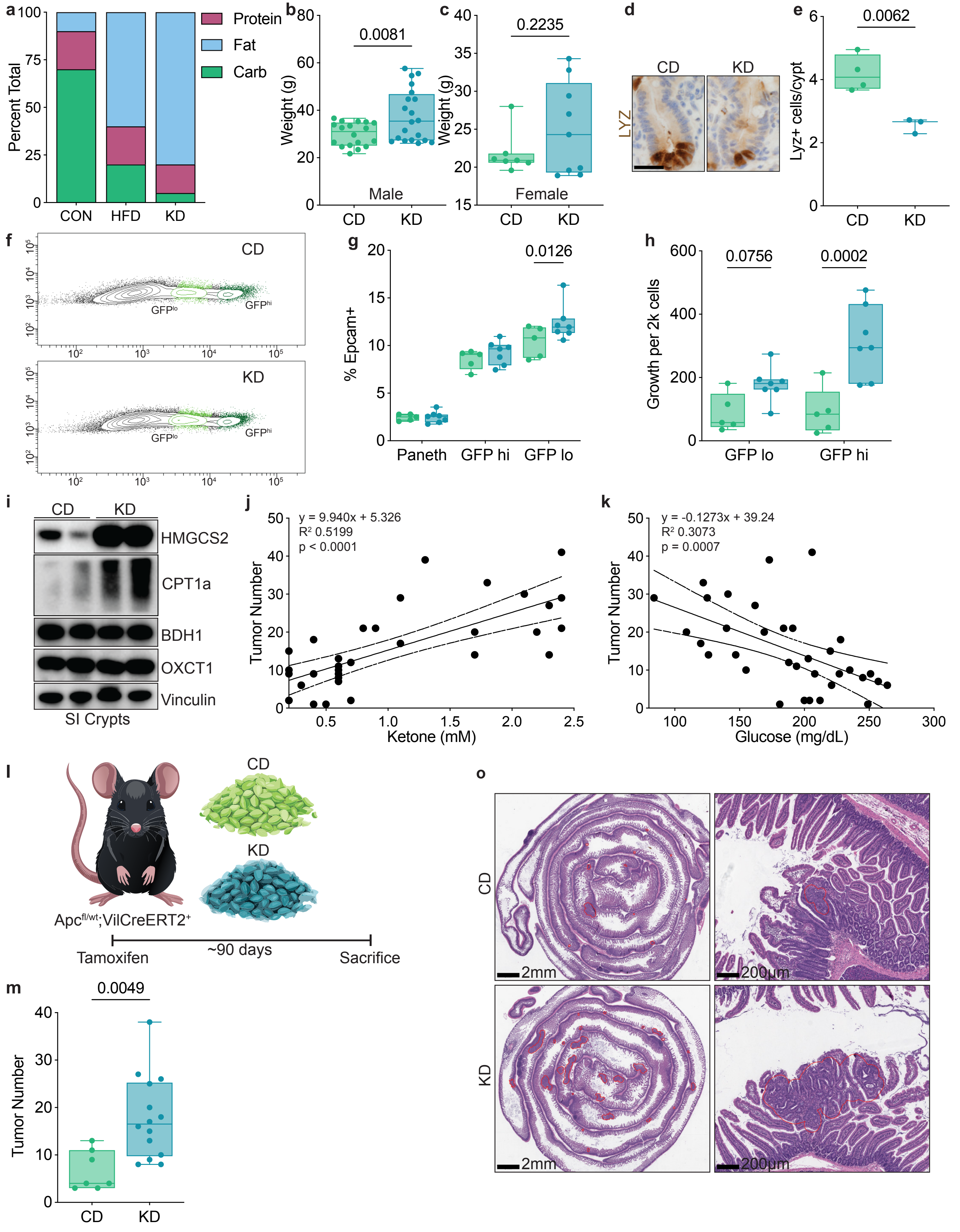

### Extended Data Figure 2

Extended Data 2

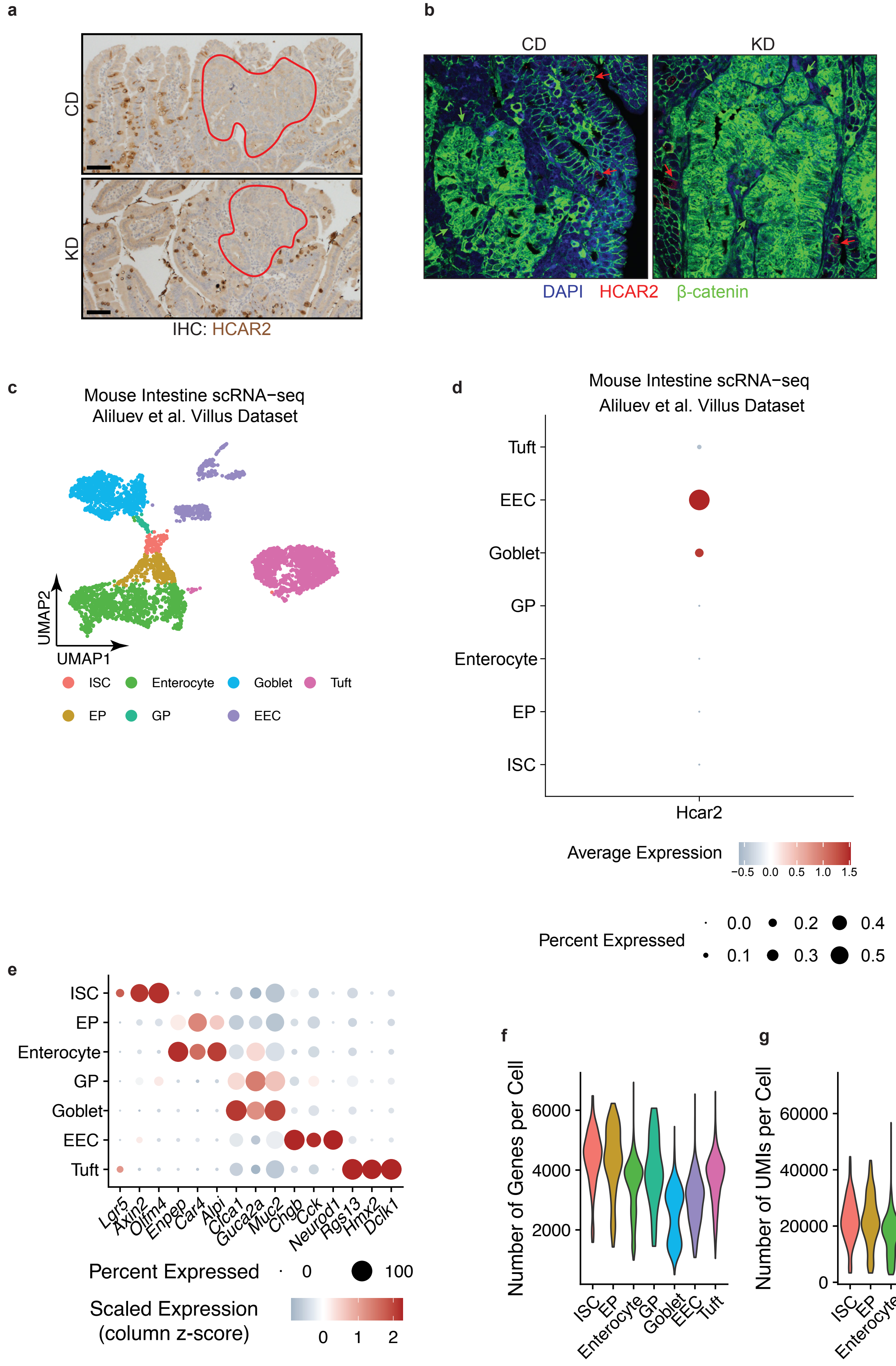

### Extended Data Figure 3

Extended Data 3

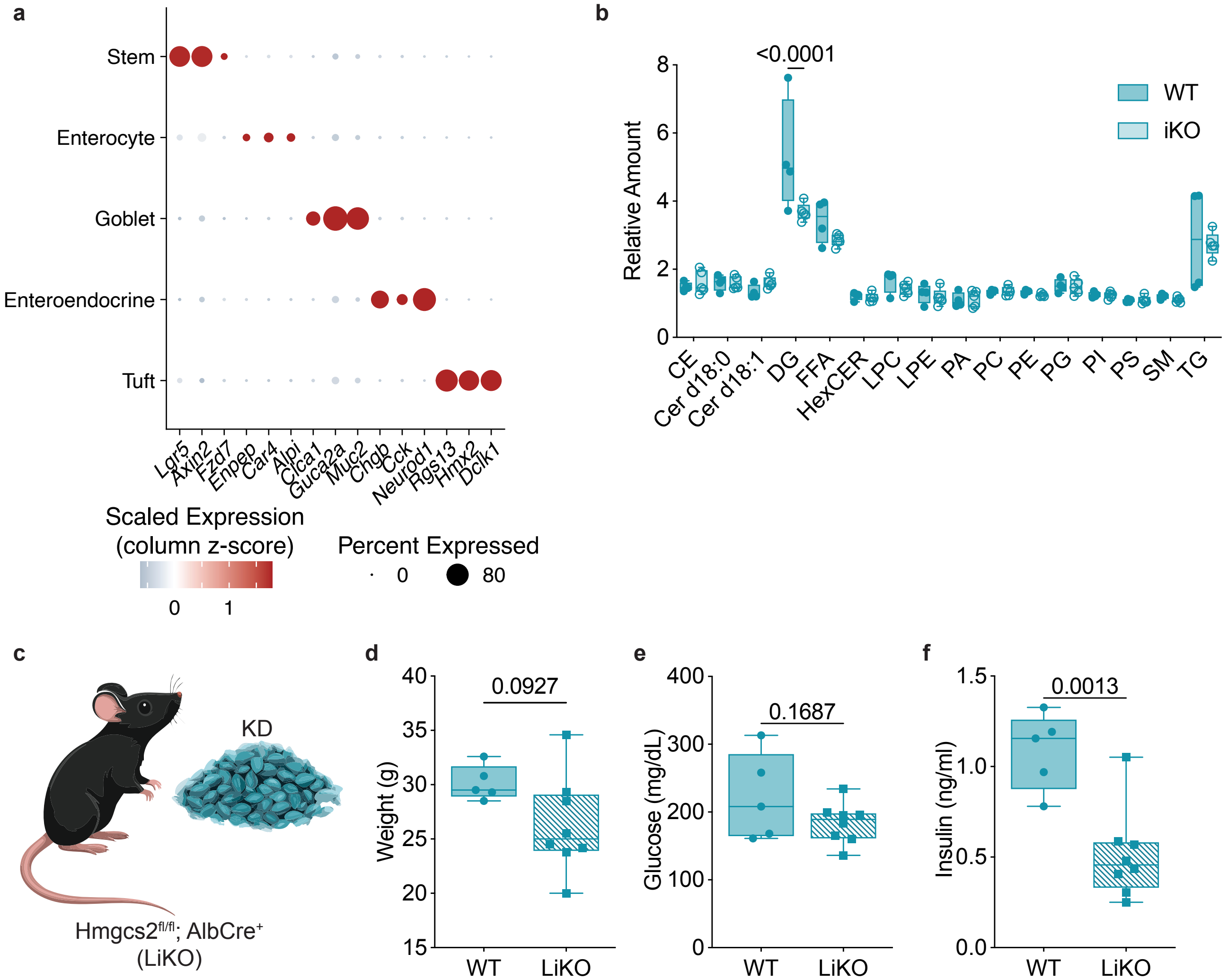

### Extended Data Figure 4

Extended Data 4

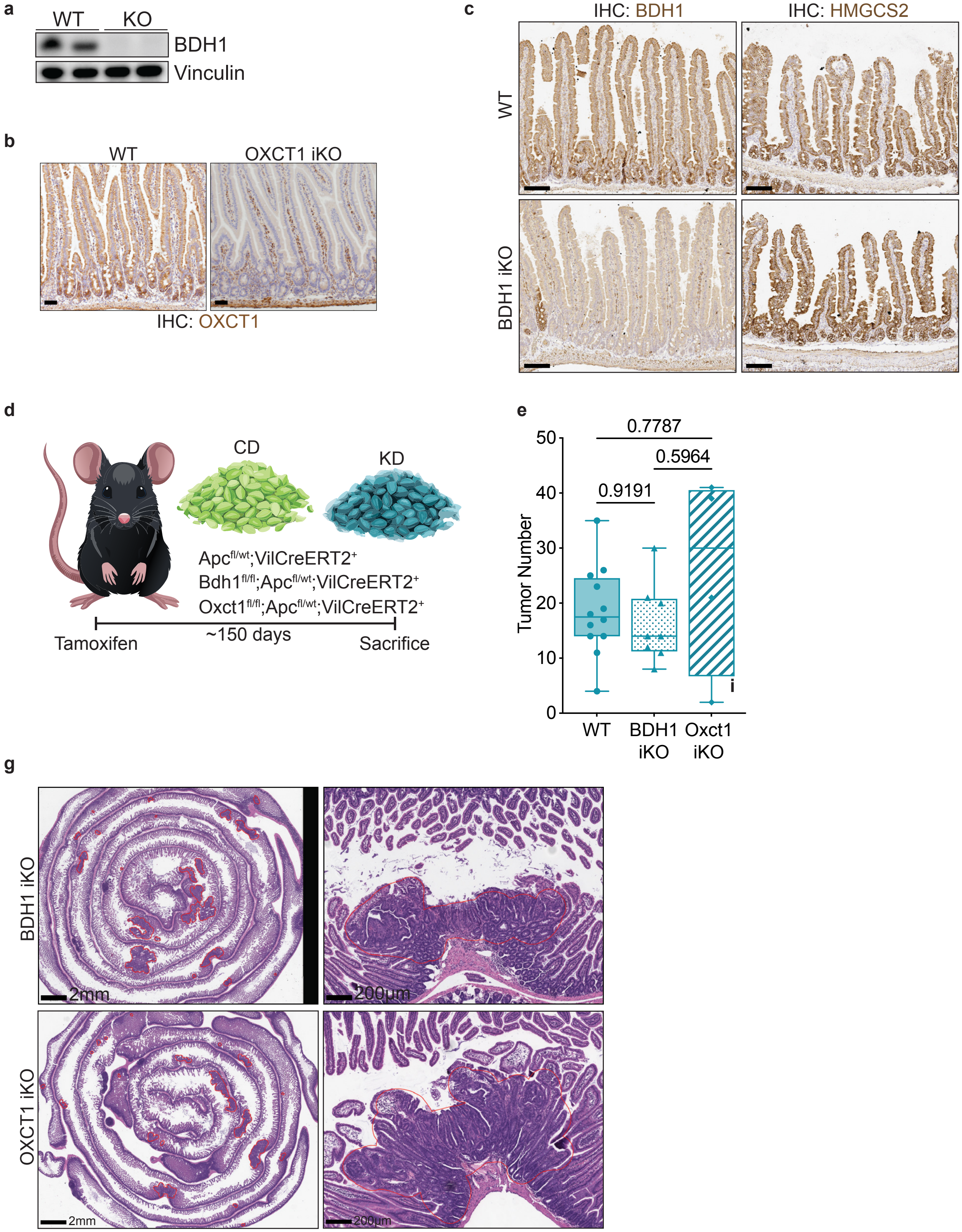

### Extended Data Figure 5

Extended Data 5

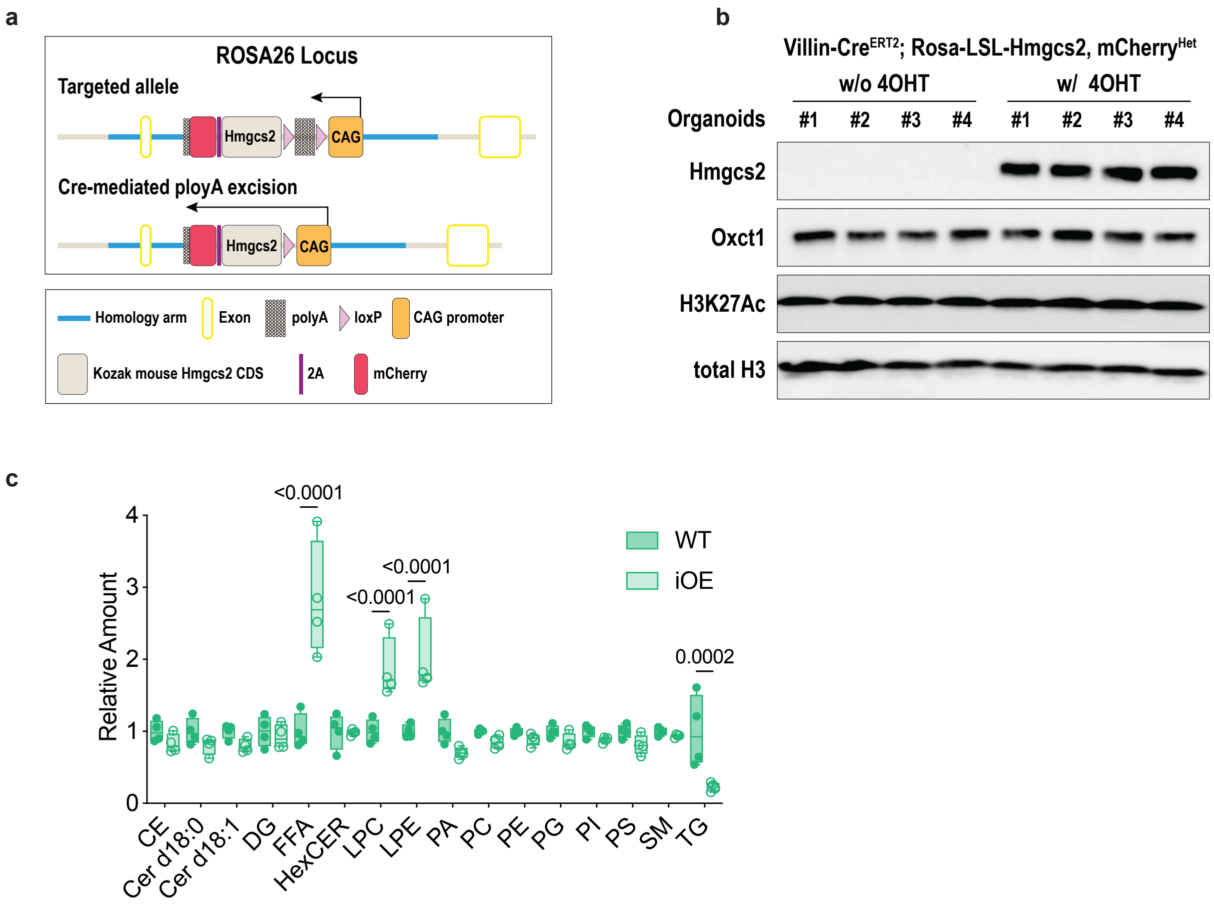

### Extended Data Figure 6

Extended Data 6

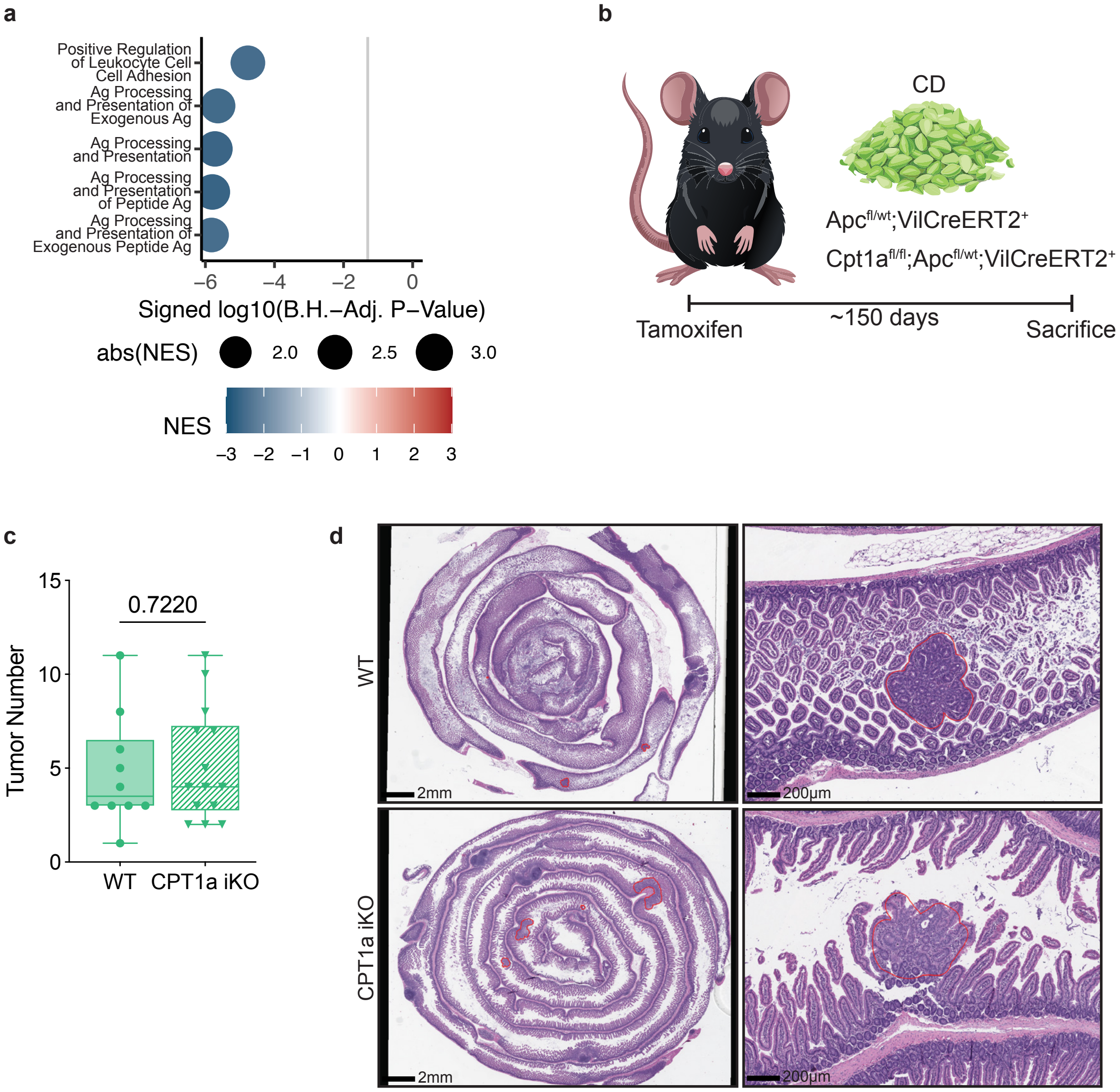

### Extended Data Figure 7

Extended Data 7

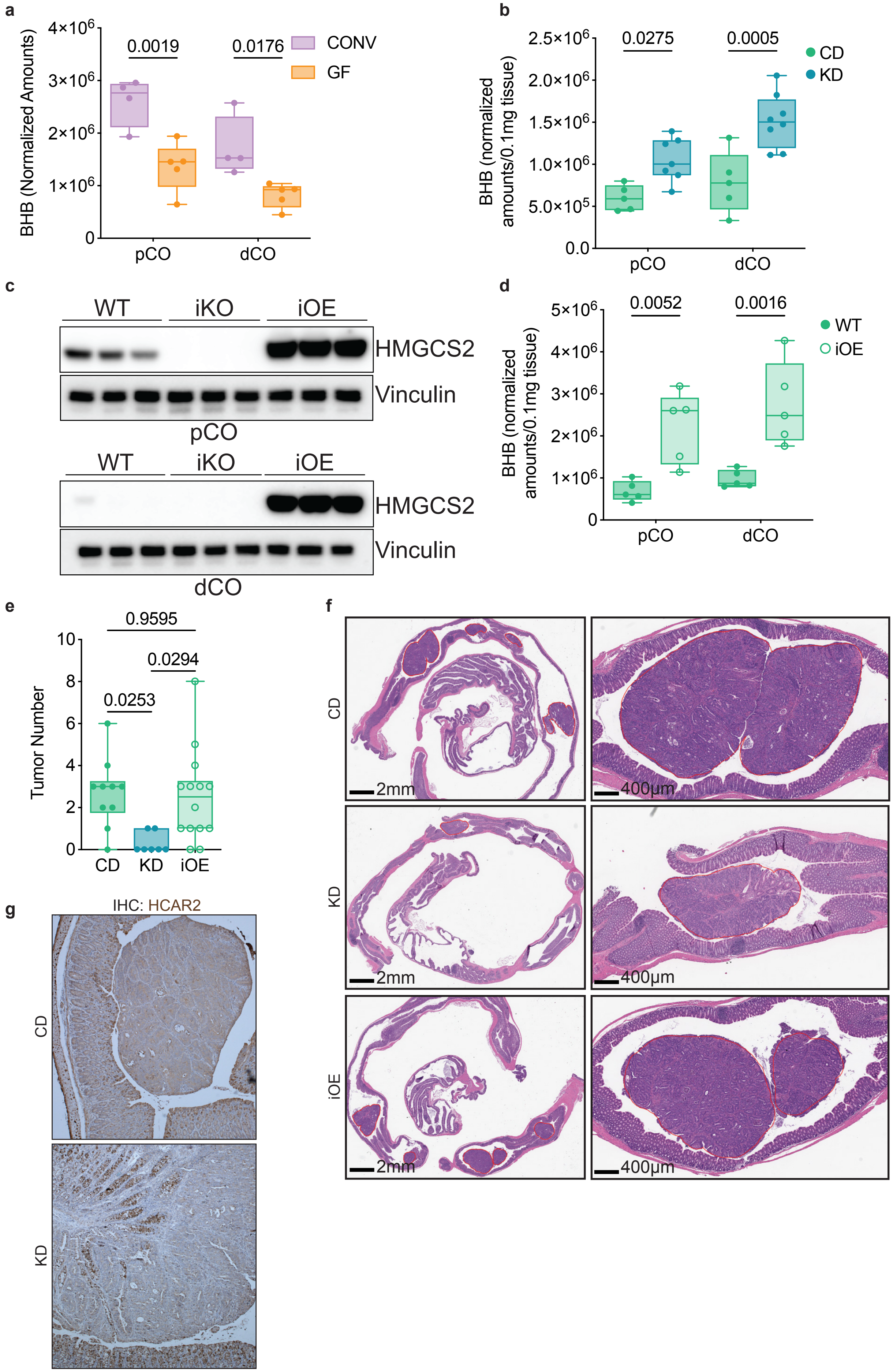
